# Distinct regions within SAP25 recruit O-linked glycosylation, DNA demethylation, and ubiquitin ligase and hydrolase activities to the Sin3/HDAC complex

**DOI:** 10.1101/2024.03.05.583553

**Authors:** Pratik Goswami, Charles A.S. Banks, Janet Thornton, Bethany Bengs, Mihaela E. Sardiu, Laurence Florens, Michael P. Washburn

## Abstract

Epigenetic control of gene expression is crucial for maintaining gene regulation. Sin3 is an evolutionarily conserved repressor protein complex mainly associated with histone deacetylase (HDAC) activity. A large number of proteins are part of Sin3/HDAC complexes, and the function of most of these members remains poorly understood. SAP25, a previously identified Sin3A associated protein of 25 kDa, has been proposed to participate in regulating gene expression programs involved in the immune response but the exact mechanism of this regulation is unclear. SAP25 is not expressed in HEK293 cells, which hence serve as a natural knockout system to decipher the molecular functions uniquely carried out by this Sin3/HDAC subunit. Using molecular, proteomic, protein engineering, and interaction network approaches, we show that SAP25 interacts with distinct enzymatic and regulatory protein complexes in addition to Sin3/HDAC. While the O-GlcNAc transferase (OGT) and the TET1 /TET2/TET3 methylcytosine dioxygenases have been previously linked to Sin3/HDAC, in HEK293 cells, these interactions were only observed in the affinity purification in which an exogenously expressed SAP25 was the bait. Additional proteins uniquely recovered from the Halo-SAP25 pull-downs included the SCF E3 ubiquitin ligase complex SKP1/FBXO3/CUL1 and the ubiquitin carboxyl-terminal hydrolase 11 (USP11), which have not been previously associated with Sin3/HDAC. Finally, we use mutational analysis to demonstrate that distinct regions of SAP25 participate in its interaction with USP11, OGT/TETs, and SCF(FBXO3).) These results suggest that SAP25 may function as an adaptor protein to coordinate the assembly of different enzymatic complexes to control Sin3/HDAC-mediated gene expression.

## Introduction

SIN3A acts as a scaffolding protein, forming a framework upon which Sin3 complexes are assembled. Sin3 co-repressors bind with histone deacetylases (HDACs), enabling Sin3-mediated transcriptional repression (1). The Sin3/HDAC family of complexes is associated with differentiation, apoptosis, proliferation, tumorigenesis and has a role in the development of cancers (2). The integrity of the Sin/HDAC complex is crucial for central nervous system development (1), methionine catabolism, and glycolysis/gluconeogenesis (3). It also plays a significant role in breast cancer metastasis (4), activating autophagy (5), contributing to pulmonary arterial hypertension (6), and influencing triple-negative breast cancer (7). In addition to the SIN3A/B paralogues, several proteins bind with the Sin/HDAC complexes including RBBP4/RBBP7 and HDAC1/2, which are shared with other transcriptional repressor complexes, and the Sin3-associated proteins SAP30/SAP30L, SUDS3, BRMS1/BRMS1L, ING1/2, ARID4A/B, SAP18, FAM60A, and SAP25 (8–11). While SIN3/HDACSin3/HDAC complexes mainly regulate gene transcription (12), several by deacetylating histone tails and altering the chromatin environment, components of the Sin3/HDAC complex interact with proteins carrying various enzymatic activities (11, 13, 14). Specifically, SIN3A has been linked to OGT, an O-linked N-acetylglucosamine transferase (12) and the TET DNA demethylases (13, 14).

First reported in 2006, TheSIN3A-associated protein of 25 kDa SAP25 was first described associated with mouse mSIN3/SIN3/HDAC complex and shown to participate in transcriptional repression (11). More recently, we showed that human SAP25 captured complexes containing the Sin3 subunits SUDS3 and SAP30 (15). SAP25 has also been associated with various additional components of epigenetic-modifying complexes, such as OGT and TET2(11, 14, 16), with an L254F-OGT variant upregulating SAP25 in X-linked intellectual disability (17). Furthermore, Tet2 purified complexes were found to contain both OGT and Sin3 complex components SAP25, SIN3A, and SAP30 (14). In addition, John and coworkers recently proposed that IFIT1 regulates gene expression programs of the immune response by coordinating the removal of Sin3 repressor complexes containing SAP25from the IFNB1, IRF7, and IRF8 promoters (18). Finally, a genome-wide DNA methylation study listed the SAP25 gene in the top 10 positions with differentially methylated CpG sites in patients with active inflammatory bowel diseases (19),while altered expression of SAP25 was observed in microglia (20), contributing to Alzheimer’s disease (21).

In this study, we focus on the SAP25 protein to elucidate its protein interaction network and role within the Sin3/HDAC complex. Mapping networks of protein interactions helps us to understand the structure of protein complexes, how they might regulate biological processes, and the impact of the aberrant function of protein complexes in different diseases (22). Affinity Purification Mass Spectrometry (AP-MS) is one of the main approaches to map protein interaction and involves using an exogenously expressed affinity-tagged bait to capture endogenous prey proteins. Proteins copurifying with the bait are identified and quantified using mass spectrometry (15, 23). AP-MS data provides significant information regarding protein-protein interactions under physiological conditions (24).

First, we conducted affinity purifications of proteins from Flp-In™-293293 cells stably expressing Halo-Tag SAP25 and analyzed purified proteins using mass spectrometry. Our analysis of proteins associated with SAP25 revealed its involvement not only with the Sin3/HDAC complex, but also with additional proteins with enzymatic function. These included the previously observed O-linked N-acetylglucosamine transferase and methylcytosine dioxygenases, but also novel associations with the SCF(FBXO3) E3 ubiquitin ligase and the ubiquitin carboxyl-terminal hydrolase 11, USP11. Next, since it had been previously observed that the C-terminal region of mouse SAP25 has an LXXLL motif required for interacting with the PAH1 domain of mSinA (11, 25), we generated a series of mutant proteins to determine whether these additional complexes could associate with SAP25 independently of Sin3/HDAC. In addition to showing that mutations within the LXXLL motif of human SAP25 disrupts its interaction with human Sin3/HDAC, we further defined additional regions of SAP25 that are necessary and sufficient for its interaction with different enzymatic complexes.

Overall, our results highlight the involvement of SAP25 as an adaptor protein capable of independently recruiting enzymatic activities to the Sin3/HDAC, potentially deploying a range of activities involved in chromatin modification and remodeling, beyond histone deacetylation. Taken together, these findings propose a broader role for SAP25 within the Sin3/HDAC complex and provide a foundation for future explorations into its function.

## Results

### SAP25 captures both the Sin3 complex and additional enzymatic and regulatory protein complexes

To determine SAP25 interacting partners, we first constructed a cell line stably expressing a recombinant version of SAP25 (NP_001335606.1) fused with an N-terminal Halo-tag (Halo-SAP25) in Flp-In™-293 cells. We used the weaker CMVd2 promoter in our Sin3 complex cell lines to express the bait proteins at a level that more closely reflects endogenous protein expression levels, even though SAP25 is not endogenously expressed in HEK293 cells (15). The resulting cell extracts were subjected to Halo affinity purification and the isolated proteins identified using Multidimensional Protein Identification Technology (MudPIT) (15, 26). We next used QPROT statistical analysis (27) to compare enrichment of proteins detected in Halo-SAP25 samples with proteins detected in controls (Fig. 1A). This analysis defined a set of 143 Halo-SAP25 enriched proteins with log_2_FC > 2 and Z statistic > 3 (Fig. 1A and Table S1A, rows 5-148).

**Figure 1:**
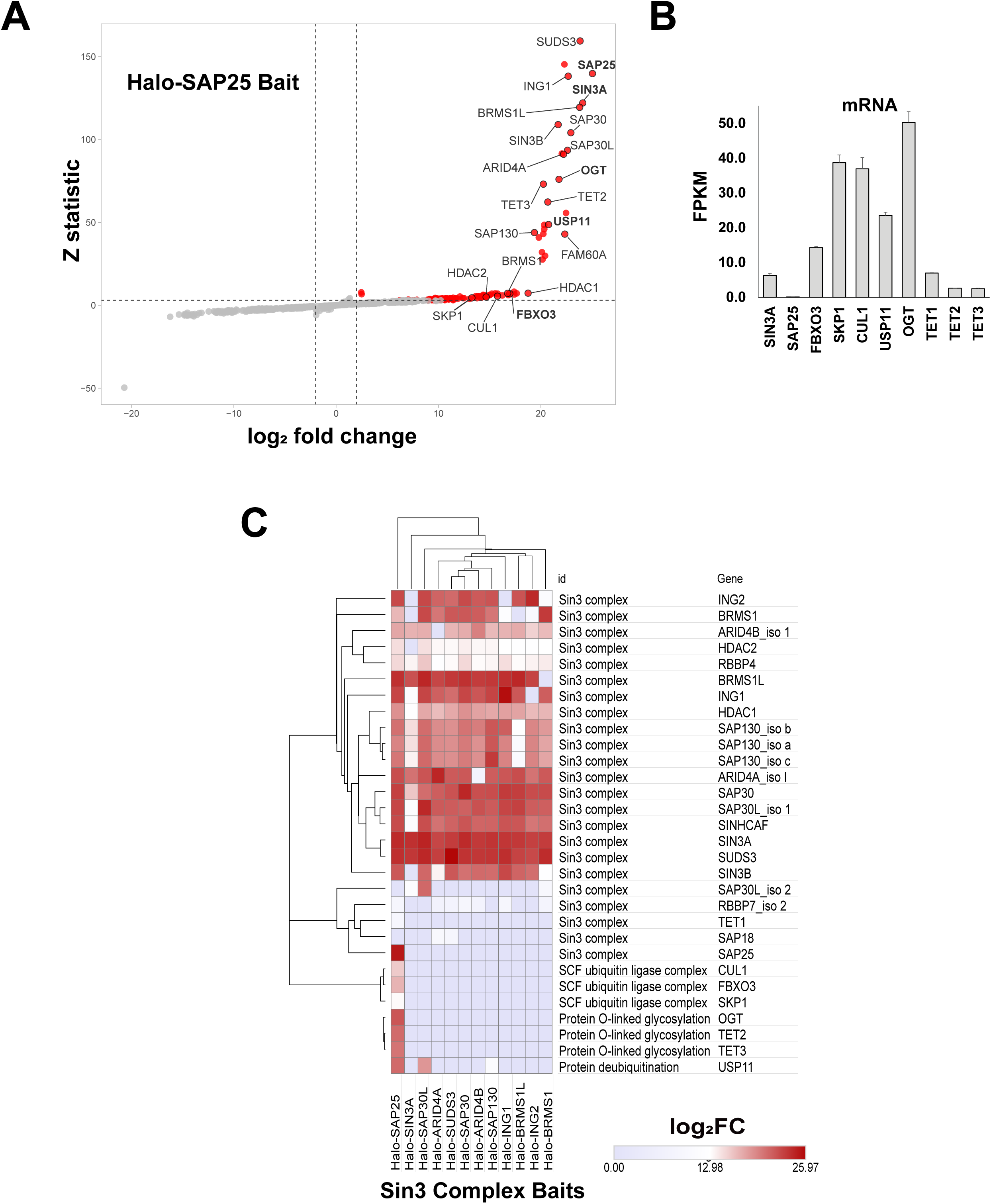
SAP25SAP25 captures a distinct subset of proteins that are not part of the Sin3 complex. (73) ***A.*** Significance plot showing proteins enriched by stably expressed Halo-SAP25 (Flp-In™-293 cells) compared with controls (log_2_FC > 2, Z statistic > 3). Significantly enriched proteins include subunits of the Sin3 complex, as well as an O-linked glycosylase (OGT), DNA demethylases (TET2/3), an E3 ubiquitin ligase complex (FBXO3, SKP1, CUL1), and ubiquitin hydrolase USP11 (Table S1A). ***B*.** Expression of SAP25 associated proteins in HEK293 cells in the absence of Halo-SAP25. FPKM for the indicated transcripts are taken from our previously published RNAseq dataset (32) available from the NCBI GEO repository under accession number GSE79656. Values are the mean of three biological replicates and error bars represent standard deviation. **C.** Hin3other enzymatic and regulatory proteins observed in purifications from Halo-SAP25 or from 1111 other stable cell lines expressing Halo-tagged Sin3 complex subunits. Values are log_2_FC (enrichment compared with Flp-In™-293 cell control purifications) (Table S1A). The comparative heatmap was generated using the Morpheus online tool (https://software.broadinstitute.org/morpheus). Clustering was performed with Euclidean distance and average linkage.

As expected, a GO term enrichment analysis of these SAP25 associated proteins using ClueGO (pV < 0.01) identified a highly enriched (pV < 0.0005) cluster of GO terms related to “Sin3-type complex” (Figure S1, pink nodes). Among the proteins identified in this cluster were the 9 Sin3 subunits we had previously identified as core Sin3 complex subunits (SIN3A, SIN3B, SUDS3, SAP30, SAP30L, HDAC1, HDAC2, RBBP4 and RBBP7) (15). In total, in addition to SAP25, we identified all 17 Sin3 complex subunits that we had previously identified copurifying with Halo-SAP30 (15) in our SAP25 purifications with relatively high abundance (dNSAF values > 0.0003) and reproducibly across biological replicates (Table S1B). This association of human SAP25 with the Sin3 complex was consistent both with a previous study in which exogenously expressed mouse SAP25 was recognized as an integral component of the mSin3 complex (11, 15), and with our previous observation by Western blotting that exogenously expressed human SAP25 captured the Sin3 subunits SUDS3 and SAP30 (15).

In addition to the Sin3 complex, we also identified the components of several protein complexes that had not previously been found to copurify with SAP25. Importantly, we identified three types of proteins with enzymatic functions: 1) components of the E3 ubiquitin ligase complex SCF(FBXO3), 2) the O-Linked N-Acetylglucosamine (GlcNAc) transferase OGT together with its established binding partners TET2 and TET3, and 3) the deubiquitinase USP11 (Fig. 1A). These proteins have previously been shown to regulate gene transcription (12, 16, 28, 29). The observation of OGT with SAP25 is also consistent with several investigations that suggest that Sin3 complex, SAP25, OGT, and TET family proteins (TET1, TET2, and TET3) collaborate in regulating transcription (12, 14, 16, 30)..

### A subset of the SAP25 interactome does not copurify with the Sin3 complex in the absence of SAP25

Next, we wanted to understand whether these complexes were copurifying with the Sin3 complex in the absence of SAP25. We had previously determined that SAP25 is not expressed in HEK293 cells (15), which hence serve as a natural “knockout” system to decipher interactions in the presence or absence of SAP25 (Fig. 1B). Therefore, we expanded our proteomic profiling and compared the Halo-SAP25 purification with purifications using other Sin3 subunits as baits. For this, we used eleven Sin3in3 complex-associated baits previously studied in our lab (31).

These include: SIN3A, SUDS3, SAP30, SAP30L, ARID4A, ARID4B, BRMS1, BRMS1L, ING1, ING2, and SAP130 (Tables S1C-M). These proteins were stably expressed in Flp-In™-293 cells with an N-terminal Halo-tag. Proteins copurifying with the Sin3 subunit baits were again identified by MudPIT, and their enrichment compared with controls (log_2_FC) calculated using the QPROT algorithm (Table S1A).

Having determined enrichment values, we compared these values for Sin3 complex subunits, SCF(FBXO3), OGT/TET, and USP11 across the different Sin3 complex baits (Fig. 1C). Specifically, we used hierarchical clustering performed with Euclidean distance as a metric and average linkage as a method. First, we observed that all baits purified Sin3 complex as we expected. Second, we found that enzymatic complexes SCF(FBXO3) and OGT/TET2/3 were not observed with any other Sin3 complex baits, suggesting that their association with SAP25 is independent of the Sin3 complex. Similarly, USP11 was also present in SAP25 purifications and absent from most other Sin3 complex purifications, although we detected USP11 with Halo-SAP30L and to a lesser extent with Halo-SAP130. Interestingly, although we had previously failed to detect SAP18 in purified Sin3 complexes (15), here we also observed that Halo-ARID4A and Halo-SUDS3 each captured modest amounts of SAP18. We next checked that by expressing Halo-SAP25 we were not simply switching on expression of the new interactors making them available for copurification with SAP25/Sin3. For this we looked at RNAseq data for HEK293 cells (in the absence of exogenous protein expression) that we had generated in a previous study (32). This confirmed that although SAP25 is not expressed in HEK293 cells, SCF(FBXO3) subunits, USP11, OGT and TET proteins are all robustly expressed (Fig 1B). Thus, they are potentially available in cells in the absence of SAP25. In conclusion, the hierarchical clustering analysis supports aa model in which SAP25associates with several complexes independently of the Sin3 complex.

### Distinct SAP25 regions are responsible for its interactions with Sin3A or with other enzymatic complexes

Having determined that a subset of proteins appeared to associate uniquely with Halo-SAP25, and not with other Sin3 subunit baits in the absence of SAP25, we hypothesized that this set of proteins might bind to one or more regions within SAP25 distinct from the region that interacts with the Sin3 complex. To investigate this, we transiently expressed different mutant versions of Halo-SAP25 in 293T cells for AP-MS analysis. Analysis of transiently expressed proteins is advantageous as it allows faster screenings of different mutants (33, 34).

First, we expressed wild type SAP25 fused to an N-terminal Halo tag in293T cells (Halo-SAP25) and tested expression by fluorescence confocal microscopy and by SDS-PAGE analysis of purified complexes (Fig. 2A). The presence of SAP25 in the nucleus was consistent with its association with Sin3 complex proteins, which are known to participate in transcription repression. In addition, we also observed a pool of Halo-SAP25 in the cytoplasm. This cytoplasmic fraction is consistent with the observation by Shiio et al.(11) of a cytoplasmic pool of recombinant mouse SAP25.

**Figure 2:**
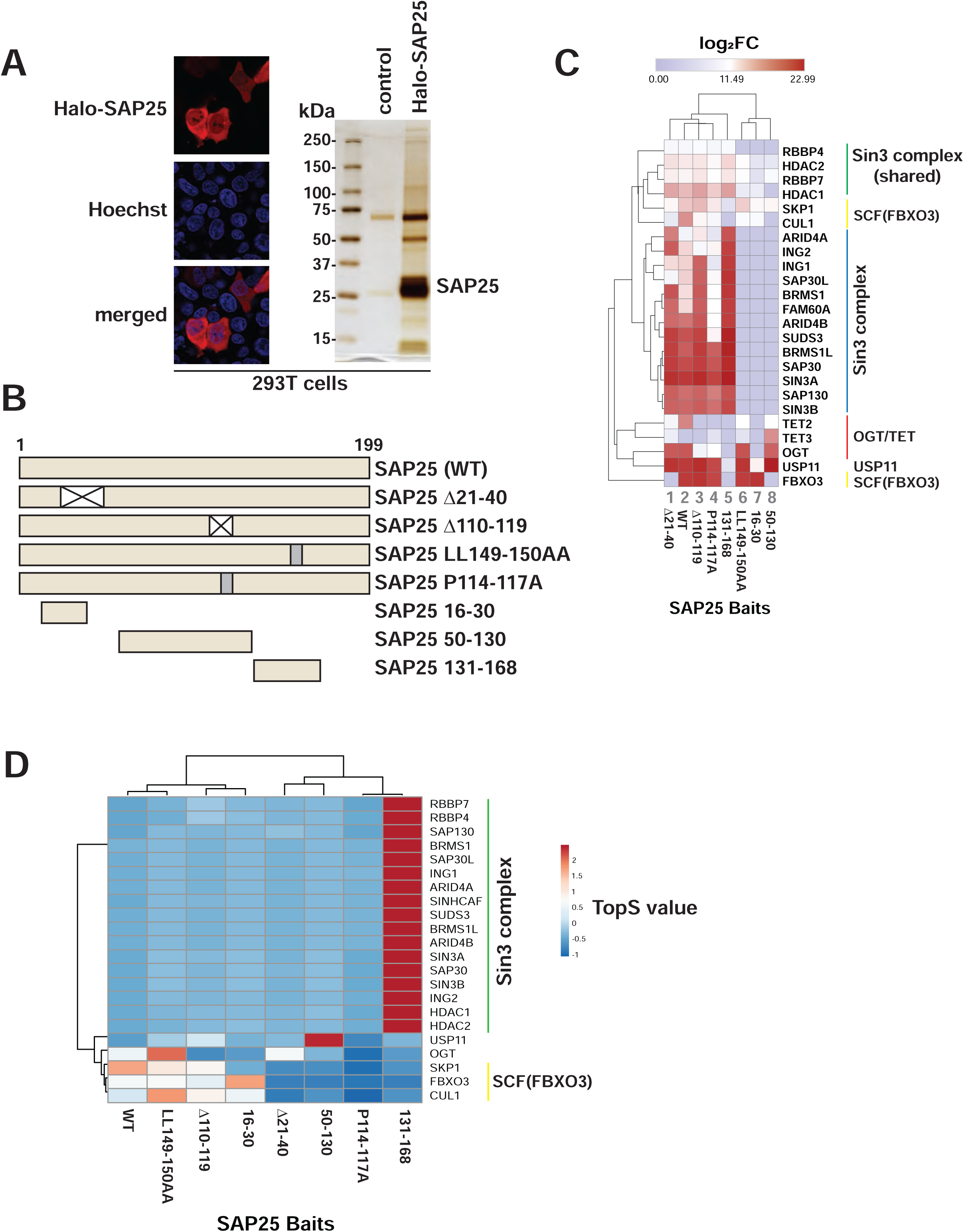
Different2Different regions of SAP25 are important for capture of either the Sin3 complex or the other enzymatic and regulatory proteins. *A.* Localization and purification of Halo-SAP25 in 293T cells. Left: Halo-SAP25 is labeled with HaloTag® TMRDirect™ fluorescent ligand (red) with DNA (nuclei) stained with Hoechst dye (blue). Right: SDSSDS-PAGE/silver stain analysis of proteins captured by Halo-SAP25 transiently expressed in 293T cells. *B.* Halo-tagged mutant versions of SAP25 were constructed as indicated for expression in 293T cells prior to AP-MS analysis. *C*. Hierarchical clustering of Sin3 complex proteins and other enzymatic and regulatory proteins observed in purifications using wild-type and mutant versions of SAP25 baits expressed in 293T cells. Values are log_2_FC with negative and missing values set to zero (Table S2A). The comparative heatmap was generated using the Morpheus online tool (https://software.broadinstitute.org/morpheus).53. Clustering was performed with Euclidean distance and average linkage. *D.* Topological analysis of SAP25 mutants. TopS scores are calculated from spectral counts assigned to proteins detected in the purifications indicated (Table S3).

Next, we asked whether different regions of SAP25 might be important for recruiting different complexes, or alternatively, whether one region in SAP25 was responsible for capturing multiple complexes. We examined the ability of mutant versions of Halo-SAP25 to enrich subunits of the different complexes compared with controls. Initially, we tested: 1) two deletion mutants (SAP25 Δ21-40 and SAP25 Δ110-119), 2) two substitution mutants (SAP25 LL149/150AA and SAP25 P114-117A), and 3) three truncation mutants covering N-terminal, middle and C-terminal regions within SAP25 (Fig. 2B). We used the QPROT algorithm (27) to calculate log_2_FC values for enriched proteins based on spectral count values (Table S2A, with the detailed lists of SAP25-associated proteins for each SAP25 construct AP-MS analysis provided in Tables S2B-I). We performed hierarchical clustering on the subset of proteins we were interested in (Fig. 2C).

First, we noticed that the components of the unique Sin3 subunits (excluding the core subunits RBBP4/7 and HDAC1/2 – which are shared with other complexes such as NuRD) clustered separately as did OGT/TET2/3 and USP11. Second, we noticed that each of our truncated versions of SAP25 enriched for a different complex: Halo-SAP25 16-30 enriches for SCF(FBXO3), but not OGT or Sin3 (Fig. 2C, lane 7); Halo-SAP25 50-130 enriches for OGT/TET and USP11 but not FBXO3 or Sin3 (Fig. 2C, lane 8); and SAP25 131-168 enriches for Sin3 subunits but not SCF(FBXO3) or OGT/TET (Fig. 2C, lane 5). Third, we noticed that our deletion mutants lost binding with components of one specific complex but not the others. For example, Halo-SAP25 Δ21-40 loses binding to FBXO3 and CUL1 but retains binding to Sin3, OGT/TET, and USP11 (Fig. 2C, lane 1). Halo-SAP25 Δ110-119 loses binding to OGT/TET but retains binding to Sin3, SCF(FBXO3), and USP11 (Fig. 2C, lane 3). Fourth, we noticed that the substitution mutant Halo-SAP25 LL149/150AA, which we generated based on a mutant of mouse SAP25 which was previously shown to lose binding to Sin3 complex (11), similarly lost binding to Sin3 complex in our experiments (Fig. 2C, lane 6). Remarkably, this mutant was still able to capture SCF(FBXO3), OGT, and USP11.

For further confirmation that the different mutant versions of SAP25 were able to capture different subsets of proteins, we also calculated TopS values (23) for Sin3 complex subunits, SCF(FBXO3), OGT/TET, and USP11 (Fig. 2D). TopS generates positive or negative scores (Table S3) for proteins based on spectral counts and these scores indicate whether the proteins are over or underrepresented in one purification compared with others (23). For the truncation mutants, we found that Sin3 complex subunits were overrepresented in Halo-SAP25 131-168 purifications (dark red segments), that USP11 was overrepresented in Halo-SAP25 50-130 purifications, and that SCF(FBXO3) was overrepresented in Halo-SAP25 16-30 purifications. In addition, we found that both SCF(FBXO3) and OGT were overrepresented in the Halo-SAP25 LL149/150AA purification, that is, their binding to SAP25 was not compromised by the substitution of these two leucine residues.

In summary, our initial analyses of these different mutant versions of Halo-SAP25 supported our hypothesis that different regions of SAP25 were likely involved in recruitment of Sin3 complex, SCF(FBXO3), USP11, and OGT/TET. Having made these observations, we next performed follow-up analyses to examine dNSAF (quantitative) values for each complex in turn and perform additional experiments to further examine the association of each complex with SAP25.

### A SAP25 C-terminal region associates with the Sin3 complex

Having determined that the region important for Sin3 recruitment was likely in the C-terminal half of SAP25, we evaluated the presence of the Sin3 complex by examining the dNSAF (abundance) values for Sin3 subunits detected in Halo-SAP25 WT, Halo-SAP25 131-168, and Halo-SAP25 LL149/150AA purifications (Fig. 3A). We found that Sin3 subunits were present in both Halo-SAP25-WT and Halo-SAP25 131-168 purifications (dark blue and light blue bars). In contrast, the majority of Sin3 subunits were not detected in Halo-SAP25 LL149-50AA purifications (Fig. 3A – absence of pink bars). Interestingly, small amounts of the subunits HDAC1/2 and RBBP4/7, which are shared among a number of HDAC complexes, were still detected in these SAP25 LL149/150AA mutant purifications. The increased abundance of Sin3 in the Halo-SAP25 131-168 compared with Halo-SAP25 WT purifications is likely due to the increase in bait abundance. The complete lists of proteins copurifying with these mutants are provided in Tables S2B, S2F, and S2H.

**Figure 3:**
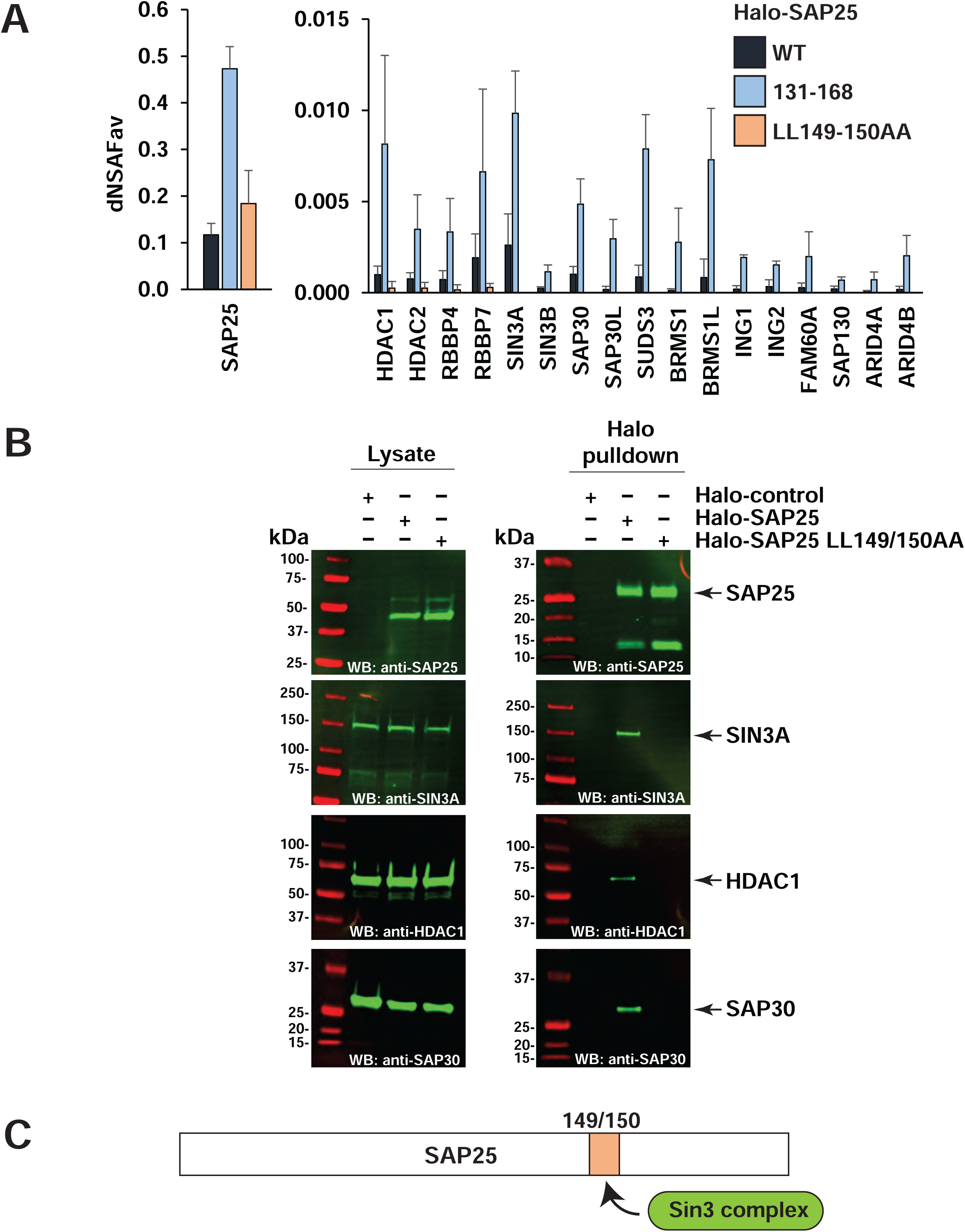
A SAP25 C-terminal region captures the Sin3 complex. ***A*.** SAP25 region 131-168 is sufficient and amino acids 149-150 are required for interaction with the Sin3 complex. Quantitative (dNSAF) values are shown for Sin3 complex proteins identified in Halo-SAP25 WT, Halo-SAP25 131-168, and Halo-SAP25 LL149-150AA purifications. Bait proteins were transiently expressed in 293T cells. Values are averages of at least three biological replicates and error bars represent standard deviation. ***B.*** Sin3 complex proteins interact with Halo-SAP25 but not Halo-SAP25 LL149-150AA. 293T cell lysates transfected with either Halo control, Halo-SAP25 or Halo-SAP25 LL149-150AA were subjected to Halo affinity capture. Copurified proteins were resolved by SDS-PAGE and identified by Western blotting using the indicated antibodies. ***C.*** Inferred position of SAP25-Sin3 complex binding. complex binding.

Next, to support our initial findings, we again expressed Halo-control, Halo-SAP25, and Halo-SAP25 LL149/150AA in293T cells and this time analyzed the eluates from the resulting purifications by SDS-PAGE followed by Western blotting. We detected Sin3 complex proteins SIN3A, HDAC1, and SAP30 as copurifying with Halo SAP25 but not with Halo-SAP25 LL149/150AA (Fig 3B -compare the last 2 lanes). This supports that the SAP25 leucine residues 149 and 150 are required for its interaction with Sin3 complex proteins (Fig. 3C).)

### The SCF(FBXO3) complex interacts with an N-terminal region of SAP25

In addition to the Sin3 complex, our AP-MS studies had indicated several potential unique interacting partners for SAP25 (Fig. 1A). Among these, we found that SAP25 captures components of the E3 ubiquitin ligase SCF(FBXO3), which promotes ubiquitination (29). SCF(FBXO3) is built from the substrate recognition protein FBXO3, the adaptor protein SKP1, the cullin CUL1, and a small (12 kDa) ring-box protein RBX1 (29). Previously, SCF(FBXO3) has been shown to target the autoimmune regulator AIRE for ubiquitination, increasing an association between AIRE and the transcription elongation factor pTEFb, and promoting proper expression of AIRE regulated genes (35). The association between SCF(FBXO3) and transcriptional regulator Sin3 was intriguing as SCF(FBXO3) had not previously copurified with any of the other Sin3 complex proteins studied by our group (Fig. 1C) (31).

To confirm that SCF(FBXO3) was recruited to SAP25 independently of the Sin3 complex, we asked which region of SAP25 might be important for the association between SCF(FBXO3) and SAP25. For this, we compared mean dNSAF values of SCF(FBXO3) complex proteins purified using the Halo tagged baits SAP25 WT, SAP25 16-30 and SAP25 Δ21-40 (Fig. 4A). We observed that both SAP25 WT and SAP25 16-30 capture three of the SCF(FBXO3) subunits, FBXO3, SKP1, and CUL1 (Fig. 4A, dark and light blue bars). We did not detect the smaller RBX1 protein. It is possible that the smaller size of this protein compromised its detection, or alternatively that SAP25 only binds SCF(FBXO3) lacking this subunit. In contrast, Halo-SAP25 Δ21-40 failed to capture the FBXO3 and CUL1 subunits of SCF(FBXO3) and captured substantially less of the SKP1 subunit, but this mutant could still capture SIN3A (Fig. 4A, pink bars). This is consistent with a model in which the SCF(FBXO3) complex binds SAP25 independently of the Sin3 complex.

**Figure 4:**
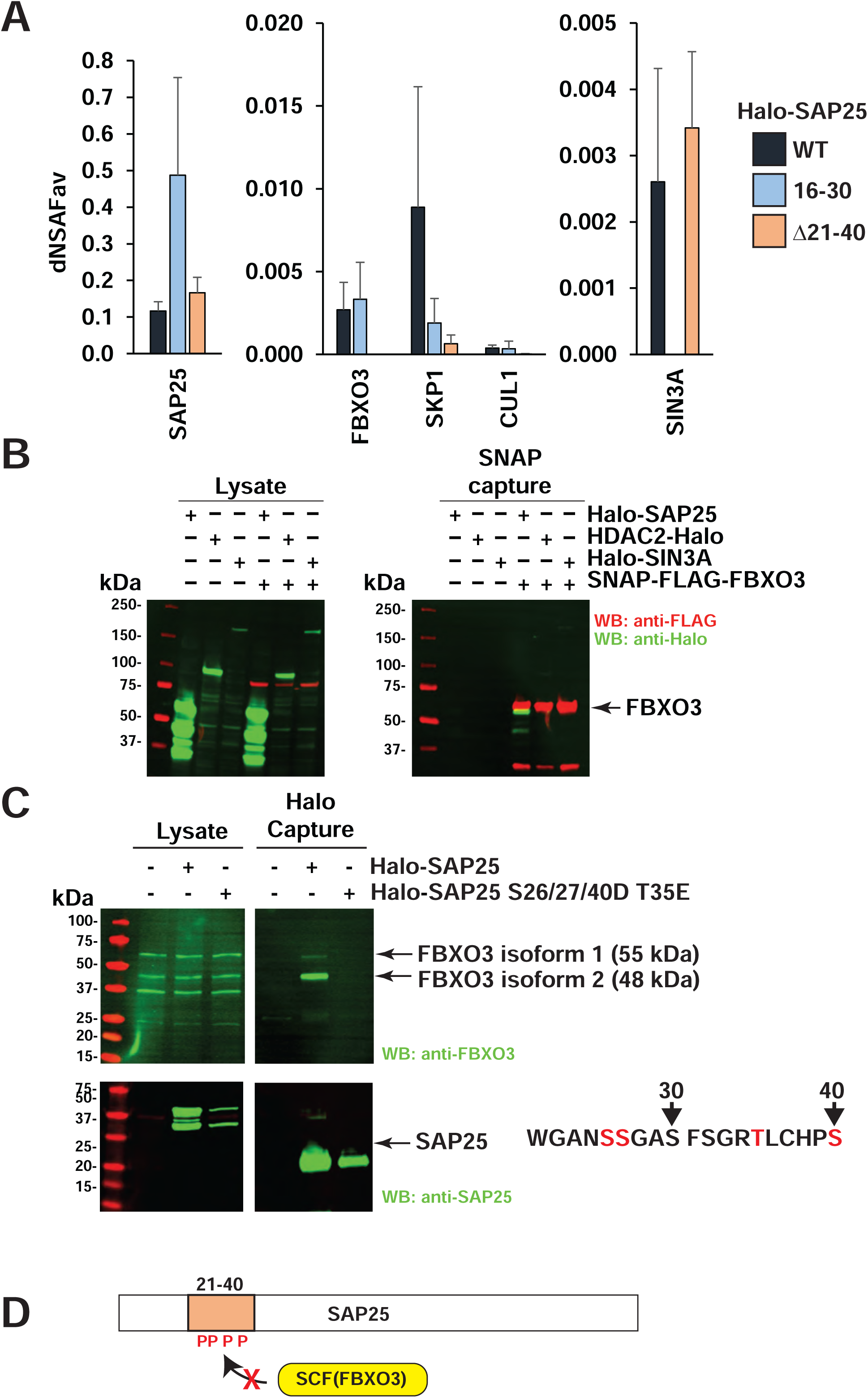
4Association between SCF(FBXO3) and an N-terminal region of SAP25. ***A.*** SAP25 region 16-30 is sufficient and region 21-40 is required for interaction with SCF(FBXO3). Quantitative (dNSAF) values for SAP25, SCF(FBXO3) complex subunits, and SIN3A are shown for Halo-SAP25 WT Halo-SAP25 16-30 and Halo-SAP25 Δ21-40purifications. Bait proteins were transiently expressed in 293T cells. Values are the average of at least three biological replicates and error bars represent standard deviation. ***B.*** FBXO3captures SAP25 but no other Sin3 complex subunits. SNAP-FLAG-FBXO3 was co-expressed with the indicated Halo-tagged proteins and the resulting lysates subjected to SNAP affinity purification. Copurified proteins were identified by SDS-PAGE followed by Western blotting using the indicated antibodies. ***C.*** Amino acid positions within SAP25 region 16-30 are important for FBXO3 capture Proteins captured by either Halo-SAP25or Halo-SAP25 S26/27/40D T35E were resolved by SDS-PAGE and detected by Western blotting using the indicated antibodies. Mutated amino acids within an N-terminal region are highlighted in red. ***D.*** Inferred position within SAP25 required for FBXO3 binding. Possible abrogation of SAP25/FBXO3 interaction by modified residues within the binding region are indicated by red Ps.

To provide additional evidence that SCF(FBXO3) does not interact directly with the Sin3 complex, we built a recombinant version of FBXO3 with an N-terminal SNAP tag and tested its ability to capture either SAP25 or Sin3 complex components SIN3A or HDAC2 (Fig 4B). We transfected 293T cells with SNAP-FLAG-FBXO3 and either Halo-SAP25, HDAC2-Halo or Halo-SIN3A and subjected the resulting cell lysates to SNAP affinity purification. After purification, as expected, we detected the presence of Halo-SAP25 in cells transfected with SNAPSNAP-FLAG-FBXO3. In contrast, SNAP-FLAG-FBXO3 could not capture HDAC2-Halo or Halo-SIN3A in the absence of SAP25 (Fig. 4B). To further support that the region including amino acids 21-40 of SAP25 is important for recruitment of FBXO3, we usedFBXO3useda Halo-SAP25 S26/27/40D T35E mutant, which contains four point-mutations within the SAP25 region 21-40. These amino acid substitutions are designed to mimic possible phosphorylation of these serine/threonine residues. The resulting Western blot analysis shows that endogenous FBXO3 copurifies with Halo-SAP25 WT but not with the Halo-SAP25 S26/27/40D T35E mutant (Fig. 4C). This supports that SAP25 amino acids within region 21-40 are important for controlling the association between SAP25 and FBXO3 (Fig. 4D).

### OGT interacts with a central region of SAP25 not required for recruitment of either the SCF(FBXO3) or Sin3 complexes

Transcription factors can be modified on serine or theonine residues by O-linked N-acetylglucosamine (O-GlcNAc) monosaccharides (36–39). Previous evidence suggests that the enzyme responsible for catalyzing this reaction, O-GlcNAc transferase (OGT), is associated with murine Sin3 complexes. Specifically, in vitro translated mSin3A was captured by GST-OGT, and HA-OGT coimmunoprecipitated mSin3A in Cos-7 cells (12).In addition, direct interaction of TET2 and TET3 with OGT has also been observed and linked with transcription regulation (16, 40). However, we had not previously detected OGT in our Sin3 purifications (Tables S1C-M) and the mechanism of recruitment of OGT to the Sin3 complex is still not fully understood.

Following our initial observation that Halo-SAP25 Δ110-119 could enrich Sin3 complex but not OGT (Fig. 2C), we considered whether OGT might be recruited to SAP25 independently of the Sin3 complex via a central region within SAP25. To examine whether SAP25 could capture OGT independently of the other complexes, we plotted dNSAF values for several representative proteins of these complexes for the SAP25 WT, SAP25 50-130, and SAP25 Δ110-119 purifications (Fig. 5A). We found that while SAP25 WT and SAP25 50-130 could both capture OGT and OGT binding partner TET2, the mutant with a disrupted central region (SAP25 Δ110-119) failed to capture OGT and TET2. The interaction between SAP25 50-130 and OGT was unlikely to be via either Sin3 or SCF(FBXO3) complexes as neither representative subunits SIN3A nor FBXO3 were detected in SAP25-50-130 purifications. Interestingly, USP11 was detected in all purifications, suggesting a different binding mechanism between SAP25 and USP11.

**Figure 5:**
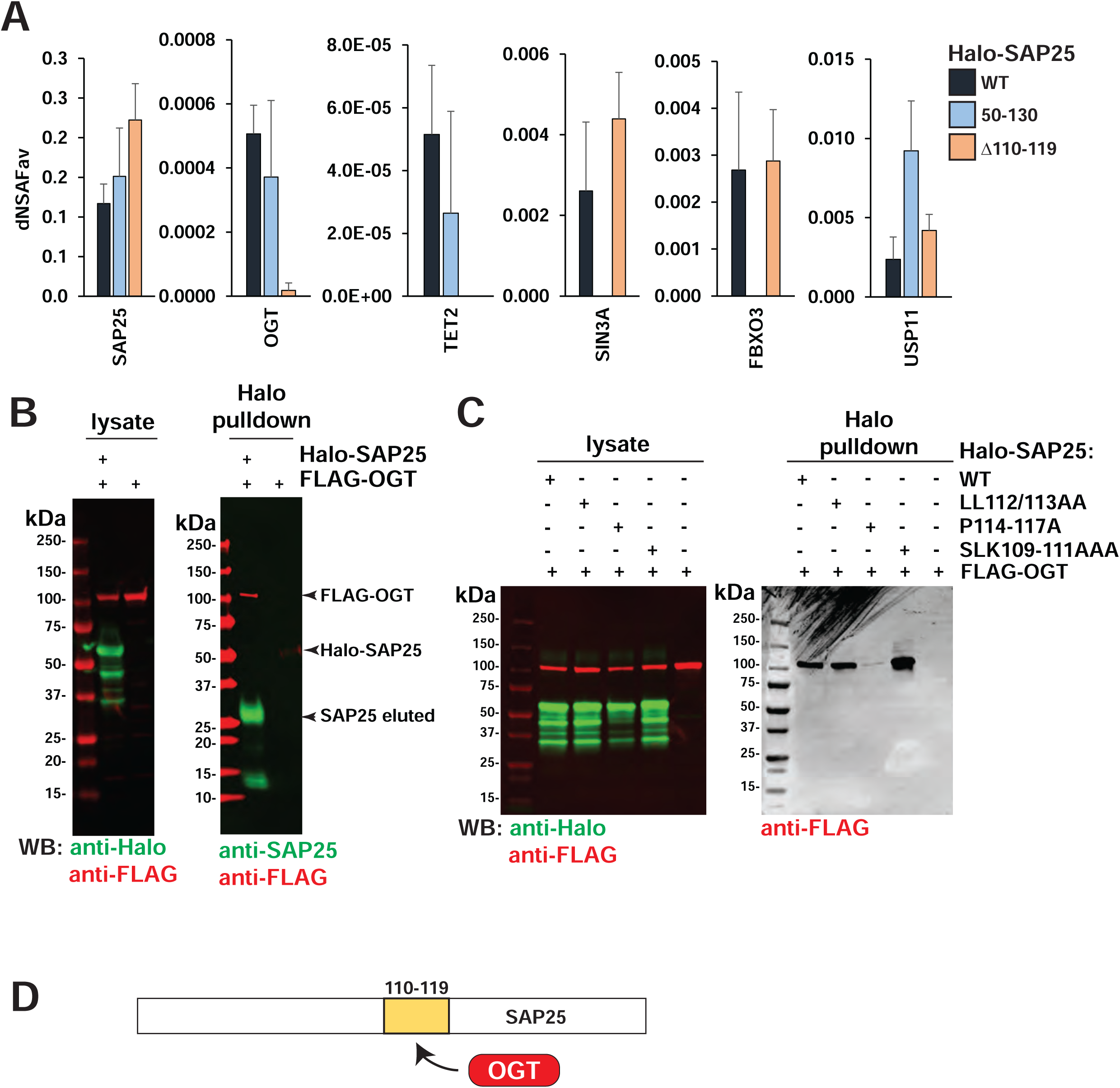
5OGT/TET interacts with a central SAP25 region. ***A.*** SAP25 region 50-130 is sufficient and region 110-119 is required for OGT/TET capture. Quantitative (dNSAF) values are shown for selected proteins identified in purifications prepared using the indicated Halo-tagged versions of SAP25. Values are averages of at least three biological replicates and error bars represent standard deviation. ***B.*** Halo-SAP25 captures FLAG-OGT. FLAG-OGT was expressed with or without Halo-SAP25 in 293T cells. Proteins identified in the resulting Halo-affinity purifications (eluted by TEV cleavage of Halo) were identified by SDS-PAGE followed by Western blot analysis using the indicated antibodies. ***C.*** Effect of disrupting sub-regions within Halo-SAP25 110-119 on OGT. FLAG-OGT was on OGT capture was co-expressed with the indicated versions of SAP25, and the resulting Halo purifications analyzed by SDS-PAGE followed by Western blotting using the indicated antibodies. ***D.*** SAP25 region required for OGT binding.

Finally, we made further mutations in the vicinity of the SAP25 110-119 region and tested the ability of two new versions of SAP25, Halo-SAP25 LL112/113AA and SLK109-111AAA to capture recombinant FLAG-OGT. Having confirmed that Halo-SAP25 WT could capture FLAG-OGT as expected (Fig. 5B), we observed that, although we captured negligible FLAG-OGT with Halo-SAP25 P114-117A (Fig. 5C), mutations just before this region (SLK109-111AAA and LL112-113AA) did not abrogate SAP25 capture of OGT. Thus, although some changes to the SAP25 110-119 region disrupt the SAP25/OGT interaction, other modest changes do not.

In conclusion, both our AP-MS data Western blot analysis support that OGT can be recruited to SAP25 independently of the Sin3 complex. We consistently identified OGT in SAP25 purifications (Fig. 1, Fig. 2C) together with the OGT binding partners TET2 and TET3 (14, 16),, but did not identify OGT copurifying with any other Sin3 complex baits, including Halo-SIN3A, in the absence of SAP25 (Fig. 1B). We mapped SAP25 region 110-119 as necessary for recruitment of OGT but not the Sin3 complex, and we determined that SAP25 50-130 is sufficient for OGT capture but not for Sin3 complex capture (Fig. 5). These results suggest that SAP25 may play a vital role in recruiting OGT to the Sin3 complex to participate in modulating post-translational modifications through O-linkage of the monosaccharide, N-acetylglucosamine (O-GlcNAc).

## Discussion

### HEK293 cells as a natural knock-out line for SAP25

We have previously published several studies to characterize the interaction network between Sin3 complex components (31)using a robust AP-MS approach combining Halo affinity purification with multidimensional protein identification technology (MudPIT). Although SAP25 had been reported as a *bona fide* component of Sin3 by Shiio et al. in 2006 (11), we were never able to detect endogenous SAP25 in Sin3 complexes isolated via other Sin3 subunits. However, exogenously expressed Halo-SAP25 did capture Sin3 components (15). RNASeq analysis revealed that SAP25 was not expressed in the HEK293 cells we used for these affinity purifications (15). Intriguingly, the O-GlcNAc transferase OGT and TET1/2/3 DNA methylases, which are known to be associated with Sin3/HDAC (12–14), were also never detected from these various purifications from Flp-In™-293 cells, except in the Halo-SAP25 dataset. Therefore, SAP25 appears to be necessary for the recruitment of these enzymatic complexes to Sin3/HDAC. To test this hypothesis further, we used 293T cells as a natural SAP25 knockout cell line to identify interaction partners uniquely pulled down by SAP25 and dissect how truncations, deletions, and point-mutations within SAP25 sequence affect these interactions without the competition for interaction from endogenous full-length SAP25.

To SAP25To identify SAP25 unique interactors, we compared the Halo-SAP25 copurified proteins with the ones identified from AP-MS analysis of eleven Halo-tagged Sin3Sin3 complex baits (Fig.1). In addition to the recovery of OGT and TET1/2/3 only in the presence of exogenous SAP25, two additional proteins/protein complexes with established enzymatic activities captured our attention: an FBXO3 based SCF ubiquitin ligase and a deubiquitinase (USP11), both of which have yet to be linked to the Sin3/HDAC complex. Having established that Halo-SAP25 captures unique interactors, we considered the mechanisms that might explain this. One possibility was that SAP25 facilitates binding of these other complexes to other Sin3 complex subunits. A second possibility was that SAP25 functions as an adaptor protein that provides a platform for the recruitment of different complexes via different regions of SAP25.

To test the possibility that SAP25 functions as an adaptor protein, we set out to map the regions of SAP25 responsible for capturing the different protein complexes. Consistent with previous observations with mouse SAP25 (11), we demonstrate that human SAP25 interacts with the Sin3 complex at via amino acids 149-150 positioned near the C-terminus of SAP25. Although we observed that this C-terminal region determines the capture of all Sin3 complex subunits (Fig. 2), it is possible that the interaction is mediated via the SIN3A scaffold subunit. Shiio and coworkers had previously established that GST-tagged mouse SAP25 could capture in vitro translated mSin3A in the absence of other Sin3 subunits. We therefore propose that human SAP25 recruits Sin3 complexes, likely via SIN3A, through a region in the SAP25 C terminus with SAP25 amino acids LL149/150 required and 131-168 sufficient for the interaction (Fig. 3).

We next investigated whether other protein complexes with enzymatic enzymatic activities dock with different regions of SAP25. We show that the N-terminus of SAP25 associates with three components of SCF(FBXO3). Deletion of AAs 21-40 results in loss of association, while expression of a much shorter SAP25 (AAs 16-30) is sufficient for SCF(FBXO3) capture (Fig. 4).

Finally, we found that a central region of SAP25 (AAs 50-130) captures both OGT/TET and USP11 protein without capturing the Sin3 or SCF(FBXO3) complexes (Fig. 5). USP11 recruitment likely does not depend on OGT/TET as deleting SAP25 amino acids 110-119 results in the loss of OGT but not USP11 (Fig. 5). Therefore, we have established a model in which distinct discrete regions of SAP25 are responsible for docking with Sin3, SCF(FBXO3), OGT/TET, and USP11 (Fig. 6). Taken together, these findings establish a distinct role for SAP25 as an adaptor protein that can separately recruit SCF(FBXO3), USP11, and OGT/TET to the theSin3 complex.

**Figure 6.**
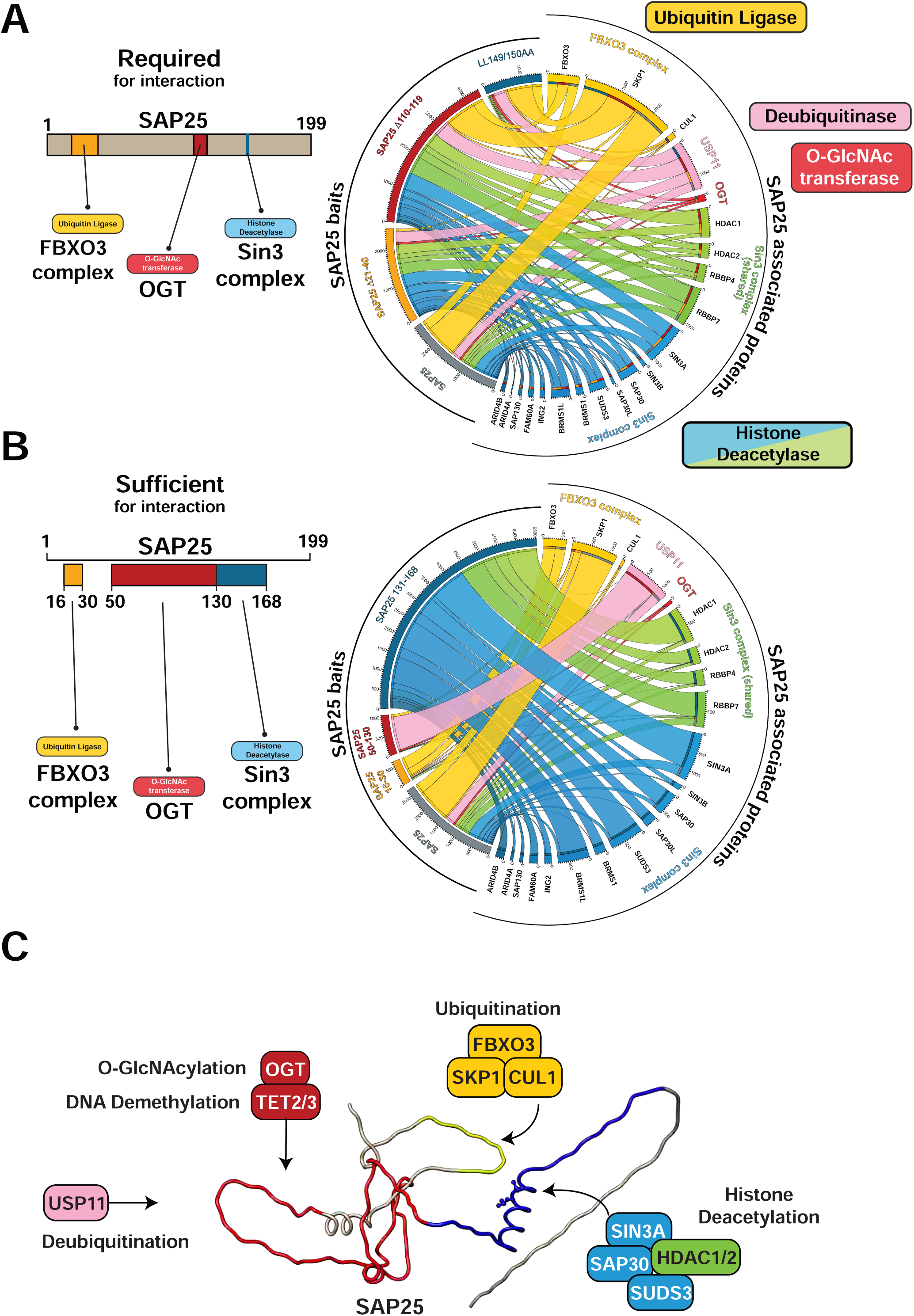
Overview of the regions of SAP25 required or sufficient for capture of SCFSCF(FBXO3), OGT, or the Sin3 complex. ***A-B.*** Quantitative comparison of Sin3 complex, SCF(FBXO3) complex, USP11, and OGT captured by the indicated versions of SAP25 expressed in 293T cells. Values on the Circos diagrams (74) are mean dNSAF x 100,000 (calculated from a minimum of three biological replicates). Subunits shared between the Sin3 and NuRD complexes are indicated by green ribbons. ***C.*** Predicted structure of SAP25 (AlphaFold AF-Q8TEE9-F1) with regions 16-30 (yellow), 50-130 (red), 131-168 (blue) and lysines 149/150 (blue atoms) indicated. indicated

Having established that SAP25 likely provides a platform for various complexes to assemble independently but in proximity, we now consider the possible roles these complexes might play while docked to SAP25. Four of these complexes are involved in protein post-translational modification – ubiquitination (SCF(FBXO3), deubiquitination (USP11), O-GlcNAcylation (OGT), or deacetylation (Sin3/HDAC) – while the TET proteins are DNA demethylases (Fig. 6). Although it is well established that the Sin3 complex represses gene transcription by deacetylating histones, the targets of the other complexes when recruited to Sin3/HDACs by SAP25 are not known.

### Association of SAP25/Sin3 with USP11

USP11 is a ubiquitin hydrolase that post-translationally modifies target proteins by removing ubiquitin. Proteins can be ubiquitinated through conjugation to either the N-terminus or to one of the seven lysines within ubiquitin. Although K48 linked ubiquitination is well known for targeting proteins for destruction by the proteosome, cleavage assays have indicated that USP11 favors targeting K6, K11, K33, and K63 linked ubiquitin chains (41). Of these, K63 ubiquitination has been well characterized and is thought to coordinate a variety of processes including DNA repair (reviewed in (42)). Timely transcriptional repression is important for initiating DNA repair and several lines of evidence have also linked USP11 to transcriptional repressor complexes.

First, USP11 was shown to physically interact with the components MTA2, MBD3, and HDAC2 of the transcriptional repressor NuRD (28). The HDAC1/2 histone deacetylases and RBBP4/7 subunits form a core complex shared between NuRD and Sin3/HDAC complexes (Fig. 6A-B – green ribbons). In the same study, NuRD and USP11 were also recruited together to UV laser induced sites of DNA damage where USP11 was shown to deubiquitinate histones H2A (K119Ub) and H2B (K120Ub). Ting and co-workers proposed that USP11 might be influencing the DNA repair process by helping to remove repair factors during the later periods of DNA damage response (28). In addition to NuRD, USP11 was found to interact with the PCGF2 (MEL18) subunit of the polycomb repressive complex PRC1 (43). PRC1 complex acts as an E3 ubiquitin ligase to ubiquitinate histone H2AK119. In this context, USP11 was not seen to affect H2A ubiquitination, but rather to target PRC1 complex subunits (MEL18, BMI1, and RING1) for deubiquitination. Maertens and coworkers proposed that here, USP11 was modulating the degradation of these PRC1 subunits, and hence the availability of PRC1 for repression of target genes such as *INK4a*. Finally, USP11 targets and stabilize the promyelocytic protein PML (44). PML interacts directly with the Sin3 complex components mSin3A and HDAC1 (45) and a pool of SAP25 has also been found colocalized with PML in PML nuclear bodies (11) consistent with a possible role for SAP25 in recruiting USP11 to the Sin3 complex for targeting Sin3-bound PML. To conclude, USP11 has been accompanying several complexes involved in transcriptional repression including NuRD, PRC1 and PML, and may target either complex subunits or histones (H2A/B) for deubiquitination.

### Association of SAP25/Sin3 with SCF(FBXO3)

In contrast to USP11, the SCF(FBXO3) complex functions as an E3 ubiquitin ligase, but like USP11, SCF(FBXO3) has also been observed to interact with PML via the FBXO3 subunit (46). Shima and coworkers proposed that SCF(FBXO3) does not target PML itself, but rather targets transcriptional coactivators (p300 and HIPK2) for ubiquitination and degradation. PML interaction with SCF(FBXO3) inhibits this degradation of coactivators, stabilizing transcription. This raises the possibility that SAP25 also controls SCF(FBXO3) targeting of coactivators. SCF(FBXO3) also targets the transcription factor AIRE vis the FBXO3 subunit (35). Shao and coworkers found that although SCF(FBXO3) does appears to target the transcription factor for degradation, ubiquitination of AIRE also increases its binding to transcription elongation factor P-TEFb, ensuring proper transcription elongation of AIRE regulated genes. Finally, SCF(FBXO3) may play a role in controlling inflammation. Specifically, FBXO3 targets another F box protein FBX12 for degradation, stimulating inflammation (47). Similarly, SAP25 has also been proposed to help control expression of inflammatory genes (18).

### Association of SAP25/Sin3 with OGT and TET proteins

OGT, an enzyme that catalyzes the addition of O-linked N-acetylglucosamine (O-GlcNAc) monosaccharides (12), is essential for cell survival (48), and is enriched in the nucleus where it plays a functional role in regulating transcription (reviewed in (49)). Nuclear OGT may target histones. Specifically, O-GlcNAcylation of histone H2B by OGT promotes its monoubiquitination at K120 (the same residue deubiquitinated by USP11) (50), possibly to create an anchor to recruit the E3 ubiquitin ligase RNF20/40.

Several lines of evidence have previously suggested that OGT can act cooperatively with the Sin3 complex. Initially, Yang et al. tested the ability of OGT to capture in vitro translated murine Sin3A and determined that OGT contacts the mSin3A PAH4 domain. By recruiting Gal4-Sin3A to promoters with or without exogenous OGT, they then demonstrated that OGT and mSin3A act synergistically to repress transcription (12). Although they found that OGT and Sin3A interact *in vitro*, the exact mechanism of OGT recruitment to the Sin3 complex in cells might be more complex as we needed SAP25 for capture of OGT together with SIN3A (Fig. 1B). Later studies also confirmed a role for murine Sin3A in functioning cooperatively with OGT. Having characterized TET1 and TET2 as OGT interactors in mouse ES cell nuclei, Vella et al. purified complexes via OGT and found them to contain mSin3A, HDAC1 and TET1/2 (51). After showing that OGT and TET colocalize at transcription start sites, with TET being required for OGT recruitment to promoters, they confirmed O-GlcNAcylation of both TET and mSin3A and proposed them as OGT targets.

Other lines of evidence have focused on the interplay between the Sin3 complex and the TET family of demethylases. Using mouse ES cells (which do express SAP25), mouse TET2 was recently found copurifying with Sin3, SAP25, and OGT, with evidence suggesting a role for TET2 (unrelated to its catalytic activity) in recruiting Sin3 complex to active enhancers (14). In contrast to mouse ES cell experiments, and in line with our observations, Chen et al (52) detected OGT but not Sin3 complex subunits copurifying with either tagged TET2 or TET3 purified from 293T cells (which we have shown lack SAP25 (15)).). Our finding that SAP25 acts as a bridge between OGT and Sin3 explains the absence of Sin3 proteins in these TET2/3 purifications. It is possible that a different recruitment mechanism governs the association of TET1 with Sin3. TET1, OGT and Sin3 complex subunits do copurify (53), but recently, TET1 was shown to interact with the SIN3A PAH1 domain (the same domain that interacts with SAP25) (13). A separate recruitment mechanism for TET1 might explain why we find TET2/3 but not TET1 copurifying with SAP25 (Fig. 1B).

### Mining the Halo-SAP25 datasets for additional unique associations

We have shown that the combination of statistical analysis with deleted/mutated/truncated versions of SAP25 yielded biologically relevant and unique interaction partners. We filtered the full-length and mutant datasets for proteins that were significantly enriched in pull-downs from cells stably expressing Halo-SAP25 WT (Table S1A), not detected with other Sin3/HDAC subunits in the absence of SAP25 (Table S1A), and significantly depleted in specific deletions or mutation and/or recovered by pull-down with the corresponding region (Table S2A). Two additional proteins passed this filtering criteria and, intriguingly, both are LIM domain proteins belonging to the zyxin/ajuba family: the Wilms tumor protein 1-interacting protein (WTIP, NP_001073905.1) and the LIM domain-containing protein ajuba isoform 1 (NP_116265.1). Both proteins reproducibly interacted with full-length Halo-SAP25 purified from stable cell lines and neither were detected in any of the 43 independent AP-MS analyses using 11 Sin3 subunits as baits (Table S1A, ranks 27-28); neither were detected in the Δ110-119 and P114-117A SAP25 mutants but both were recovered from the corresponding affinity purification using region 50-130 as bait (Table S2A, ranks 27-28), identifying this region of SAP25 as sufficient for this interaction. WTIP (54) and Ajuba (55) contain 3 LIM domains in their C-terminal region and have been shown to interact with each other and another LIM domain protein LIMD1 (56), which was detected at low levels in the Halo-SAP25 transient affinity-purifications and exhibited the same pattern of depletion/recovery in the SAP25 mutant analyses (Table S2A, ranks 29), which suggests that Halo-SAP25 pulled them down as a complex.

LIM domains are made of two zing fingers loops separated by a short hydrophobic linker of two amino acids (as reviewed in (57)). LIM domain proteins are considered scaffolds in the assembly of multiple complexes. As such, WTIP, Ajuba, and LIMD1 have been involved in many cellular processes such as cell proliferation (58–61) and fate determination (59); cytoskeletal organization (62, 63); gene silencing through microRNA (56) and repression of gene transcription (64–67); mitosis (63, 68); cell differentiation (66) and migration (69). One of the main characteristics of LIM domain proteins of the zyxin/ajuba family is their ability to shuffle between the cytoplasm and the nucleus (59, 64), hence carrying signaling cues from external stimuli and sites of cell-cell adhesion on the plasma membrane to the nucleus where they function as co-repressors of specific genes. In this context, their association with SAP25 is intriguing because of the existence of a pool of cytosolic SAP25 (Fig. 2A). Another interesting observation is that the nuclear localization of xyzin, Ajuba orthologue in mouse, requires its O-GlcNAcylation by OGT (70), another protein we have shown here to interact with the same middle region of SAP25. Finally, Ajuba has been linked previously to histone deacetylase activity by co-immunoprecipitation and size exclusion chromatography of nuclear extracts from Jurkat T-cells followed by Western blots. Because of the antibody-based approach to characterize these interactions, whether specific HDAC1/2/3-containing complexes were involved could not be determined then. Since SAP25 is highly expressed in Jurkat cells (clone E6-1; from (71)), SAP25 could serve as the link between Ajuba and Sin3 complexes containing HDAC1/2.

### Conclusions

Our work reveals the interplay between SAP25, several complexes with enzymatic and regulatory functions, and the Sin3 complex. We hypothesize that SAP25 is acting as an adaptor protein and recruits these complexes to the Sin3 complex, possibly to regulate transcription by modulating post-translational and DNA modifications. Evidence from previous studies suggests that SAP25 may act as a docking protein as it does not display any recognizable DNA-binding motifs (11). Our topological and comparative proteomics analysis of Sin3 complex baits supports our hypothesis. Recently, SAP25 and HDAC were linked with the expression of inflammation and antiviral response genes (18). Furthermore, FBXO3, a distinctive interacting partner of SAP25, is associated with breast cancer metastasis induced by PI3K and is associated with USP4 stabilization (72). The presence of USP11 in SAP25 purifications could indicate a role for SAP25 in coordinating transcriptional repression by Sin3/HDAC with DNA damage repair. These results suggest promising directions for further research into the function of SAP25, which could help to determine whether SAP25 functions to recruit these enzymatic activities to the Sin3 complex to regulate transcriptional repression.

## Acknowledgements

The research presented here was supported by the Stowers Institute for Medical Research and the National Institute of General Medical Sciences of the National Institutes of and R35GM145240 (M.P.W.). The content is solely the responsibility of the authors and does not necessarily represent the official views of the National Institutes of Health.

## Author Contributions

Conceptualization: CAB, PG, MPW

Investigation: PG, CAB, BB, MS

Formal Analysis: PG, CAB, BB, MS, LF

Resources: CAB, JT

Data Curation: CAB, PG, BB, MS, LF

Writing: PG, CAB, LF, MPW

Supervision: LF, MPW

Funding acquisition: MPW

## Declaration of Interests

The authors declare that they have no conflicts of interest related to the work presented here.

## Materials and Methods

### Materials

Magne® HaloTag® beads (G7281), HaloTag® polyclonal antibody (G9281) and TMRDirect™ fluorescent ligand (G2991) were purchased from Promega. AcTEV protease (#12575015) was from Thermo Fisher Scientific. Rabbit anti-SAP30 (ab125187) and rabbit anti-SIN3A (ab3479) polyclonal antibodies were from Abcam (Cambridge, United Kingdom). SNAP-Capture magnetic beads (S9145S) were from NEB (Ipswich, MA). Salt Active Nuclease (SAN) was from ArcticZymes (Tromso, Norway). Rabbit anti-FBXO3 (HPA002467) was from Atlas Antibodies. Rabbit anti-HDAC1 (10197-1-AP) was from Proteintech. Rabbit anti-SAP25 (HPA062610-100UL), and mouse anti-FLAG (F3165-.2MG) were from Sigma. IRDye® 800CW labeled goat anti-Rabbit (926–3211) and IRDye® 680LT labeled goat anti-Mouse (926–68020) secondary antibodies were from LI-COR Biosciences. FLAG-OGT in pcDNA5FRT (75) was a gift from the laboratory of Joan and Ron Conaway.

### Cloning sequences to express Halo affinity tagged versions of SAP25 and Sin3 complex subunits

A codon optimized synthetic sequence (SAP25 in pIDTSmart (Amp)) coding for a 199 aa sequence matching SAP25 (originally annotated as NP_001162153.1) was used to amplify various sequences coding for different versions of SAP25 using the primers listed in Supplementary Data (cloning of full-length SAP25 in pFN21A was described previously (15)). This 199 AA sequence also matches the more recently annotated version NP_001335606.1 (SAP25 isoform 3). For transient expression, the resulting PCR products were subcloned into either pFN21A (Promega) or Halo pcDNA5/FRT PacI PmeI (described previously (76)). The construction of all stable cell lines except for Halo-SIN3A and Halo-SAP25 has been described previously (15, 31). Halo-SIN3A was cloned from ORF Clone # FHC11647 (Promega) by first mutating the codon for amino acid 109 from GCT (A) to GTT (V) to match the translated sequence to NP_056292. The resulting sequence was then cloned into CMVd2 pcDNA5 PacI PmeI (described previously (77)) using the primers listed in Supplementary Data. Stable cell line construction is described in Banks et al. (2018) (15). Halo-SAP25 was also cloned into CMVd2 pcDNA5 PacI PmeI for construction of a stable cell line. For expression of FLAG-OGT, the mutations in the OGT1 insert (N764D and K1010E) in the original plasmid were corrected using site directed mutagenesis. For expression of FBXO3, an FBXO3(NM_012175) ORF Clone was purchased from GenScript (Cat. #OHU25332D), and the open reading frame amplified using the primers listed in Supplementary Data. The resulting PCR product was subcloned into SNAP-FLAG-pcDNA5 (15).

### Preparation of whole cell extracts

For transient expression of mutant versions of SAP25, whole cell lysates were prepared from approximately 1 x 10^7^ 293T cells transiently transfected with the constructs indicated in the figure legends as described previously (76). Cells were harvested 48 hours after transfection and washed twice with ice-cold PBS. Cell pellets were frozen, thawed, and resuspended in approximately 300 μl lysis buffer containing 50 mM Tris-HCl (pH 7.5), 150 mM NaCl, 1% Triton® X-100, 0.1% sodium deoxycholate, 0.1 mM benzamidine HCl, 55 μM phenanthroline, 10 μM bestatin, 20 μM leupeptin, 5 μM pepstatin A and 1 mM PMSF. The resulting lysate was passed through a 26-gauge needle 10 times and centrifuged at 21, 000 x g for 30 minutes at 4°C. The supernatant was diluted with 700 μl of TBS (25 mM Tris-HCl pH 7.4, 137 mM NaCl, 2.7 mM KCl). Extracts were prepared from Flp-In™-293 stable cell lines as described previously (76).

### Purification of recombinant protein complexes

Lysates were incubated overnight at 4°C with Magne® HaloTag® beads prepared from 0.1 ml bead slurry (transiently transfected cells) or 0.2 ml of bead slurry (stable cell lines). The beads were isolated using a DynaMag-2 Magnet and the supernatant discarded. Beads were washed 4 times in buffer containing TBS (25 mM Tris-HCl (pH 7.4), 137 mM NaCl, 2.7 mM KCl) and 0.05% Nonidet® P40. To elute bound proteins, beads were incubated in 100 µl elution buffer (25 mM Tris-HCl (pH 8.0), 0.5 mM EDTA, 1 mM DTT, and 2 Units AcTEV) for 2h at 25°C. To purify SNAP-tagged proteins, cell lysates were allowed to incubate overnight at 4 °C with SNAP-Capture magnetic beads followed by washing with wash buffer as described above. Elution of bound proteins was accomplished by incubating the beads with a buffer containing 50 mm Tris·HCl pH 8.0, 0.5 mm EDTA, 1 mm DTT, and 1 Unit of PreScission Protease at 4 °C overnight.

### Digestion of proteins for mass spectrometry

HaloTag® purified protein complexes were TCA precipitated and resuspended in buffer containing 100 mM Tris-HCl, pH 8.5 and 8 M urea. Disulfide bonds were reduced with 5 mM TCEP (tris (2-carboxyethyl) phosphine) for 30 minutes at room temperature and bond reformation prevented by alkylation with 10 mM chloroacetamide for 30 minutes at room temperature in the dark. Proteins were initially digested with the addition of 0.1 μg of Lys-C followed by incubation in a ThermoMixer® (Eppendorf) for 6 hours at 37°C with shaking. Samples were diluted to 2 M urea by the addition of an appropriate volume of 100 mM Tris-HCl, pH 8.5. Proteins were further digested with the addition of 2 mM CaCl_2_ and 0.5 µg trypsin, and subsequent incubation in a ThermoMixer® at 37°C overnight. Reactions were quenched with the addition of formic acid (5% final concentration) prior to mass spectrometry analysis.

### Liquid chromatography mass spectrometry analysis

Digested peptides were analyzed using an LTQ Linear Ion Trap mass spectrometer (Thermo Scientific) run in positive ion mode connected to an Agilent 1100 HPLC system. Samples were first loaded offline onto 3-phase microcapillary MudPIT columns as described previously (78) and eluted into the mass spectrometer using a series of 10 ∼2-hour MudPIT steps. Chromatography gradients used a combination of Buffer A (5% acetonitrile, 0.1% formic acid), Buffer B (80% acetonitrile, 0.1% formic acid), and Buffer C (500 mM ammonium acetate, 5% acetonitrile, 0.1% formic acid).

### Processing of tandem mass spectrometry data

Resulting .raw files were converted to .ms2 files using RAWDistiller v. 1.0. The ProLuCID algorithm version 1.3.5 was used for searching. Spectra from stable cell line experiments were matched to a database containing 48080 human protein sequences downloaded from the National Center for Biotechnology Information (NCBI) (2022-04-12 release), together with 427 common contaminant sequences and shuffled versions of all sequences to estimate false discovery rates (FDRs). The total number of sequences searched was 97014. Spectra from transient transfection experiments were searched with the same database, but with the addition of 8 sequences for recombinant mutant versions of SAP25 (together with 8 further shuffled sequences). Data were searched for peptides with a static modification of 57.0215 on cysteine residues (carbamidomethylation) and a variable modification of 15.9949 on methionine residues (oxidation). A mass tolerance of 500 ppm was used for both precursor and fragment ions. Only fully tryptic peptides were considered.

The entire in-house software suite (kite), used for the processing and analysis of the mass spectrometry datasets, is available in Zenodo (79). We used the in-house software program swallow in combination with DTASelect v. 1.9 (80) to filter peptide spectrum matches (PSMs) resulting in mean spectrum, peptide, and protein FDRs below 5% (Tables S1B-M and S2B-I). The mean spectral FDR for the 36 MudPIT runs (transient transfection experiments) was 0.714% ± 0.263% (standard deviation), and the mean protein FDR was 3.958% ± 1.838% (standard deviation). For stable cell line experiments, the mean spectral FDR for the 54 MudPIT runs was 0.468% ± 0.173% (standard deviation), and the mean protein FDR was 2.133% ± 1.139% (standard deviation). The minimum peptide length was 7 amino acids. Proteins identified in different runs were compared using sandmartin (79), Contrast (80), and NSAF7 software. The parsimony option in Contrast was used to remove proteins that were subsets of others.

### Experimental design and statistical rationale

AP-MS analyses required at least three biological replicates (Tables S1-S2). The number of biological replicates for analyses involving a stable cell line was: 5 (Halo-control); 5 (Halo-SAP25). For analyses using transient cells the number of biological replicates employed was: 9 (Halo-control); 4 (Halo-SAP25); 4 (Halo-SAP25 Δ21-40); 4 (Halo-SAP25 Δ110-119); 3 (Halo-SAP25 P114-117A); 3 (Halo-Sap25 LL149-150AA); 3 (Halo-SAP25 16-30); 3 (Halo-SAP25 50-130); 3 (Halo-SAP25 131-168). We employed QPROT, for analyzing differential protein expression which is based on protein intensity data (27). A high-confidence interaction was confirmed when the interaction parameters satisfied the conditions of log_2_FC > 2 and Z-statistic > 3.

### Fluorescence microscopy

HEK293T cells were seeded to 40% confluency in MatTek culture dishes and incubated at 37 °C in 5% CO_2_ for 24 hours. Cells were then subsequently transfected with plasmid expressing Halo-SAP25. HaloTag® TMRDirect™ was introduced to the culture medium at a concentration of 20 nm to label Halo-tagged proteins. The cells were then incubated overnight at 37 °C in 5% CO_2_. Next day, cells were stained with Hoechst dye for 1 hour to label nuclei followed by washing with Opti-MEM® reduced serum medium. The cells were then visualized using an LSM-700 Falcon confocal microscope.

### Topological scoring

Topological scoring was performed for each protein as described previously (23, 81). Proteins significantly enriched with log_2_FC > 2 and Z-statistics > 3 were selected as input to the TopS Shiny application, available at https://github.com/WashburnLab/Topological-score-TopS. This application uses the mean spectral counts for each bait across all baits to derive TopS score, signifying a probability of binding. Proteins were filtered out based on TopS value greater than 10 (Table S3).

### Data availability

We have reported some of the raw mass spectrometry data used in this study previously (all searches are new to this study). A summary of the runs used here, together with details of where they were first reported is provided in Table S4. Mass spectrometry data has been deposited in the MassIVE repository (http://massive.ucsd.edu) with the data identifiers MSV000093553 (transient transfection data) and MSV000093576 (stable cell line data). Original data underlying this manuscript generated at the Stowers Institute can be accessed after publication from the Stowers Original Data Repository at http://www.stowers.org/research/publications/libpb-2438.

## Notes

### Competing Interest Statement

The authors have declared no competing interest.

