## Supplemental Data and Figures for "Distinct regions within SAP25 recruit O-linked glycosylation, DNA demethylation, and ubiquitin ligase and hydrolase activities to the Sin3/HDAC complex"

### Description of Supplementary Tables:

| | |
|---|---|
| **Table S1** | **AP-MS data for the proteins Halo-affinity purified from Flp-In™-293 cell lines stably expressing SAP25 & 11 other SIN3/HDAC subunits as baits** |
| S1A | Average dNSAF & QPROT FC/Z-statistics for proteins quantified across 12 Halo-APs of SIN3/HDAC subunits and Halo-Controls |
| S1B. | Detailed peptide counts, spectral counts, sequence coverage, and summary statistics for the proteins affinity-purified in replicate Halo-SAP25 AP-MS analyses and Halo-controls |
| S1C. | Detailed peptide counts, spectral counts, sequence coverage, and summary statistics for the proteins affinity-purified in replicate Halo-SIN3A AP-MS analyses and Halo-controls |
| S1D. | Detailed peptide counts, spectral counts, sequence coverage, and summary statistics for the proteins affinity-purified in replicate Halo-SUDS3 AP-MS analyses and Halo-controls |
| S1E. | Detailed peptide counts, spectral counts, sequence coverage, and summary statistics for the proteins affinity-purified in replicate Halo-SAP30 AP-MS analyses and Halo-controls |
| S1F. | Detailed peptide counts, spectral counts, sequence coverage, and summary statistics for the proteins affinity-purified in replicate Halo-SAP30L AP-MS analyses and Halo-controls |
| S1G. | Detailed peptide counts, spectral counts, sequence coverage, and summary statistics for the proteins affinity-purified in replicate Halo-ARID4A AP-MS analyses and Halo-controls |
| S1H. | Detailed peptide counts, spectral counts, sequence coverage, and summary statistics for the proteins affinity-purified in replicate Halo-ARID4B AP-MS analyses and Halo-controls |
| S1I. | Detailed peptide counts, spectral counts, sequence coverage, and summary statistics for the proteins affinity-purified in replicate Halo-BRMS1 AP-MS analyses and Halo-controls |
| S1J. | Detailed peptide counts, spectral counts, sequence coverage, and summary statistics for the proteins affinity-purified in replicate Halo-BRMS1L AP-MS analyses and Halo-controls |
| S1K. | Detailed peptide counts, spectral counts, sequence coverage, and summary statistics for the proteins affinity-purified in replicate Halo-ING1 AP-MS analyses and Halo-controls |
| S1L. | Detailed peptide counts, spectral counts, sequence coverage, and summary statistics for the proteins affinity-purified in replicate Halo-ING2 AP-MS analyses and Halo-controls |
| S1M. | Detailed peptide counts, spectral counts, sequence coverage, and summary statistics for the proteins affinity-purified in replicate Halo-SAP130 AP-MS analyses and Halo-controls |

| | |
|---|---|
| **Table S2** | **AP-MS data for the proteins Halo-affinity purified from 293T cell lines transiently expressing full-length SAP25 isoform 3 (NP_001335606.1 ) & 7 deleted/mutated/truncated SAP25 constructs as baits** |
| S2A | Average dNSAF & QPROT FC/Z-statistics for proteins quantified across 8 Halo-APs of SAP25 constructs and Halo-Controls |
| S2B. | Detailed peptide counts, spectral counts, sequence coverage, and summary statistics for the proteins affinity-purified in replicate Halo-SAP25 WT AP-MS analyses and Halo-controls |
| S2C. | Detailed peptide counts, spectral counts, sequence coverage, and summary statistics for the proteins affinity-purified in replicate Halo-SAP25 Δ21-40 (D5) AP-MS analyses and Halo-controls |
| S2D. | Detailed peptide counts, spectral counts, sequence coverage, and summary statistics for the proteins affinity-purified in replicate Halo-SAP25 Δ110-119 (D3) AP-MS analyses and Halo-controls |
| S2E. | Detailed peptide counts, spectral counts, sequence coverage, and summary statistics for the proteins affinity-purified in replicate Halo-SAP25 P114-117A (M10) AP-MS analyses and Halo-controls |
| S2F. | Detailed peptide counts, spectral counts, sequence coverage, and summary statistics for the proteins affinity-purified in replicate Halo-SAP25 LL149-150AA (M7) AP-MS analyses and Halo-controls |
| S2G. | Detailed peptide counts, spectral counts, sequence coverage, and summary statistics for the proteins affinity-purified in replicate Halo-SAP25 16-30 (T19) AP-MS analyses and Halo-controls |
| S2H. | Detailed peptide counts, spectral counts, sequence coverage, and summary statistics for the proteins affinity-purified in replicate Halo-SAP25 50-130 (T3) AP-MS analyses and Halo-controls |
| S2I. | Detailed peptide counts, spectral counts, sequence coverage, and summary statistics for the proteins affinity-purified in replicate Halo-SAP25 131-168 (T43) AP-MS analyses and Halo-controls |

**Table S3 Output_TopS_Transient_mutations**

S3A. TopS_values

S3B. TopS_sort

S3C. TopS_sort_highlighted

S3D. FOR FIGURE

**Table S4 Summary of AP-MS datasets used in this study**

S4A. Stable Cell Line MS runs

S4B. Transient Transfection MS runs

### Supplementary Data

#### 1. Primers used to construct vectors expressing Halo tagged versions of SAP25

a) For cloning Halo-SAP25 in pFN21A, CMVd2 pcDNA5 PacI PmeI, and SNAP-FLAG-pcDNA5:

SgfI SAP25 F 5'- GACGCGATCGCCATGACGCCTCTCGCACCCTG - 3'

PmeI SAP25 R (with stop) 5'- GACGTTTAAACTTAGGGGCAGTGAGTGTCTGGAG - 3'

b) For cloning Halo-SAP25 Δ21-40 in pFN21A:

First round PCR

N-terminal PCR product

SgfI SAP25 F 5'- GACGCGATCGCCATGACGCCTCTCGCACCCTG - 3'

SAP25 D21-40 R 5'- CAGTGGCCAAAACACAGGTCGGGGCCCG - 3'

C-terminal PCR product

SAP25 D21-40 F 5'- CCGACCTGTGTTTTGGCCACTGTACGAGGCCG - 3'

PmeI SAP25 R 5'- GACGTTTAAACTTAGGGGCAGTGAGTGTCTGGAG - 3'

Second round PCR used SgfI SAP25 F & PmeI SAP25 R primers with first round PCR products as template.

c) For cloning Halo-SAP25 Δ110-119 in pFN21A:

First round PCR

N-terminal PCR product

SgfI SAP25 F 5'- GACGCGATCGCCATGACGCCTCTCGCACCCTG - 3'

SAP25 D110-119 R 5'- CTCACAGCAAACGACGGGGAAGCCAGCATCTGG - 3'

C-terminal PCR product

SAP25 D110-119 F 5'- ATGATGGGAAGCTCCGCAAGGGTGCTGCCAC - 3'

PmeI SAP25 R 5'- GACGTTTAAACTTAGGGGCAGTGAGTGTCTGGAG - 3'

Second round PCR used SgfI SAP25 F & PmeI SAP25 R primers with first round PCR products as template.

d) For cloning Halo-SAP25 PPPP114-117AAAA in pFN21A:

First round PCR

N-terminal PCR product

SgfI SAP25 F 5'- GACGCGATCGCCATGACGCCTCTCGCACCCTG - 3'

SAP25 PPPP/AAAA R 5'- CATGATGGCAGCCGCTGCAAGCAATTT
CAAGCTTCCCATCATCTGCTGA - 3'

C-terminal PCR product

SAP25 PPPP/AAAA F 5'- GCTTGCAGCGGCTGCCATCATGTCCGCA
AGGGTGC - 3'

PmeI SAP25 R 5'- GACGTTTAAACTTAGGGGCAGTGAGTGTCTGGAG - 3'

Second round PCR used SgfI SAP25 F & PmeI SAP25 R primers with first round PCR products as template.

e) For cloning Halo-SAP25 LL149/150AA in pFN21A:

First round PCR

N-terminal PCR product

SgfI SAP25 F 5'- GACGCGATCGCCATGACGCCTCTCGCACCCTG - 3'

SAP25 LL149/150AA R 5'- GGCTCATCTGAGCAGCGCCCGTCAGGGCAAT
GAG - 3'

C-terminal PCR product

SAP25 LL149/150AA F 5'- GACGGGCGCTGCTCAGATGAGCCAGGGCGAAC - 3'

PmeI SAP25 R 5'- GACGTTTAAACTTAGGGGCAGTGAGTGTCTGGAG - 3'

Second round PCR used SgfI SAP25 F & PmeI SAP25 R primers with first round PCR products as template.

f) For cloning Halo-SAP25 16-30 in pFN21A:

SgfI SAP25 16 F 5'- GACGCGATCGCCATGGGGCCCCGACCTGTGTG - 3'

PmeI SAP25 30 R (with stop) 5'- GACGTTTAAACTCAGCTTGCGCCAGAGCTGCAG - 3'

g) For cloning Halo-SAP25 50-130 in Halo pcDNA5/FRT PacI PmeI:

SgfI SAP25 50 F 5'- GACGCGATCGCCATGGGCAGGGGGCTTAGGCC - 3'

PmeI SAP25 130 R (with stop) 5'- GACGTTTAAACTTAGGAAGGGCTAGGGCGTGG - 3'

h) For cloning Halo-SAP25 131-168 in pFN21A:

SgfI SAP25 131 F 5'- GACGCGATCGCCATGAGGGGGCCAAGCACGG - 3'

PmeI SAP25 168 R (with stop) 5'- CTGGTTTAAACTTAATCAGGGGGTCCGACAGCG - 3'

i) For cloning Halo-SAP25 LL112/113AA in pFN21A:

First round PCR

N-terminal PCR product

SgfI SAP25 F 5'- GACGCGATCGCCATGACGCCTCTCGCACCCTG - 3'

SAP25 LL112/113AA R 5'-GGCGGTGGAGCAGCTTTCAAGCTTCCCATCAT

CTGCTGAG - 3'

C-terminal PCR product

SAP25 LL112/113AA F 5'- GGAAGCTTGAAAGCTGCTCCACCGCCTC

CCATCATG - 3'

PmeI SAP25 R 5'- GACGTTTAAACTTAGGGGCAGTGAGTGTCTGGAG - 3'

Second round PCR used SgfI SAP25 F & PmeI SAP25 R primers with first round PCR products as template.

j) For cloning Halo-SAP25 SLK109-111AAA in pFN21A:

First round PCR

N-terminal PCR product

SgfI SAP25 F 5'- GACGCGATCGCCATGACGCCTCTCGCACCCTG - 3'

SAP25 SLK/AAA R 5'- AAGCAATGCCGCGGCTCCCATCATCTG

CTGAGGAG - 3'

C-terminal PCR product

SAP25 SLK/AAA F 5'- GGGAGCCGCGGCATTGCTTCCACCGCC - 3'

PmeI SAP25 R 5'- GACGTTTAAACTTAGGGGCAGTGAGTGTCTGGAG - 3'

Second round PCR used SgfI SAP25 F & PmeI SAP25 R primers with first round PCR products as template.

k) For cloning Halo-SAP25 S26/27/40D T35E in pFN21A:

First round PCR

N-terminal PCR product

SgfI SAP25 F 5’- GACGCGATCGCCATGACGCCTCTCGCACCCTG - 3’

SAP25 S26/27/40D T35E R 5’- CCAAAAGTCTGGGTGACAGAGCTCGCGGCCG
GAAAAGCTTGCGCCATCGTCGCAGTTTGC - 3’

C-terminal PCR product

SAP25 S26/27/40D T35E F 5’- GCAAACTGCGACGATGGCGCAAGCTTTTCCG
GCCGCGAGCTCTGTCACCCAGACTTTTGG - 3’

PmeI SAP25 R 5’- GACGTTTAAACTTAGGGGCAGTGAGTGTCTGGAG - 3’

Second round PCR used SgfI SAP25 F & PmeI SAP25 R primers with first round PCR products as template.

**2. Synthetic sequence in pIDTSmart used to subclone SAP25**

5’ - GCGATCGCCATGACGCCTCTCGCACCCTGGGACCCCAAATACGAAGCGAAAGCCGGGCCCCGACCT
GTGTGGGGGGCAAACTGCAGCTCTGGCGCAAGCTTTTCCGGCCGCACCCTCTGTCACCCATCATTTTGGC
CACTGTACGAGGCCGCGAGCGGCAGGGGGCTTAGGCCAGTTGCTCCAGCTACCGGCCATTGGAACGGC
CAGCAGGCACCACCAGATGCTGGCTTCCCCGTCGTTTGCTGTGAGGATGTATTTCTTAGTGACCCCCTCC
TCCCAAGGGGTCAGAGAGTTCCACTGTACCTGTCTAAAGCTCCTCAGCAGATGATGGGAAGCTTGAAAT
TGCTTCCACCGCCTCCCATCATGTCCGCAAGGGTGCTGCCACGCCCTAGCCCTTCCAGGGGGCCAAGCA
CGGCCTGGCTTAGCGGGCCCGAGCTCATTGCCCTGACGGGCCTTCTGCAGATGAGCCAGGGCGAACCCA
GACCTTCTAGCAGCGCTGTCGGACCCCCTGATCACACTAGCGATCCCCCCAGTCCCTGTGGGAGCCCTTC
CTCATCTCAAGGAGCAGATCTTTCTCTGCCACAGACTCCAGACACTCACTGCCCCTAAGTTTAAAC - 3’

**3. Primers used to construct Halo tagged SIN3A in CMVd2 pcDNA5 PacI PmeI**

SgfI SIN3A F 5’- CAG GCGATCGCC ATG AAG CGG CGT TTG GAT GAC C - 3’

PmeI SIN3A R (with stop) 5’- CAG GTTTAAACTTA AGG GGC TTT GAA TAC TGT GCC GTATTT G - 3’

**4. Primers used to construct SNAP-FLAG tagged FBXO3 in pcDNA5**

SgfI FBXO3 F 5’- CAG GCGATCGCC ATG GCG GCC ATG GAG ACC - 3’

PmeI FBXO3 R 5’- CAG GTTTAAAC CTA AAA AAG GCG TGA GCA GCG G - 3’

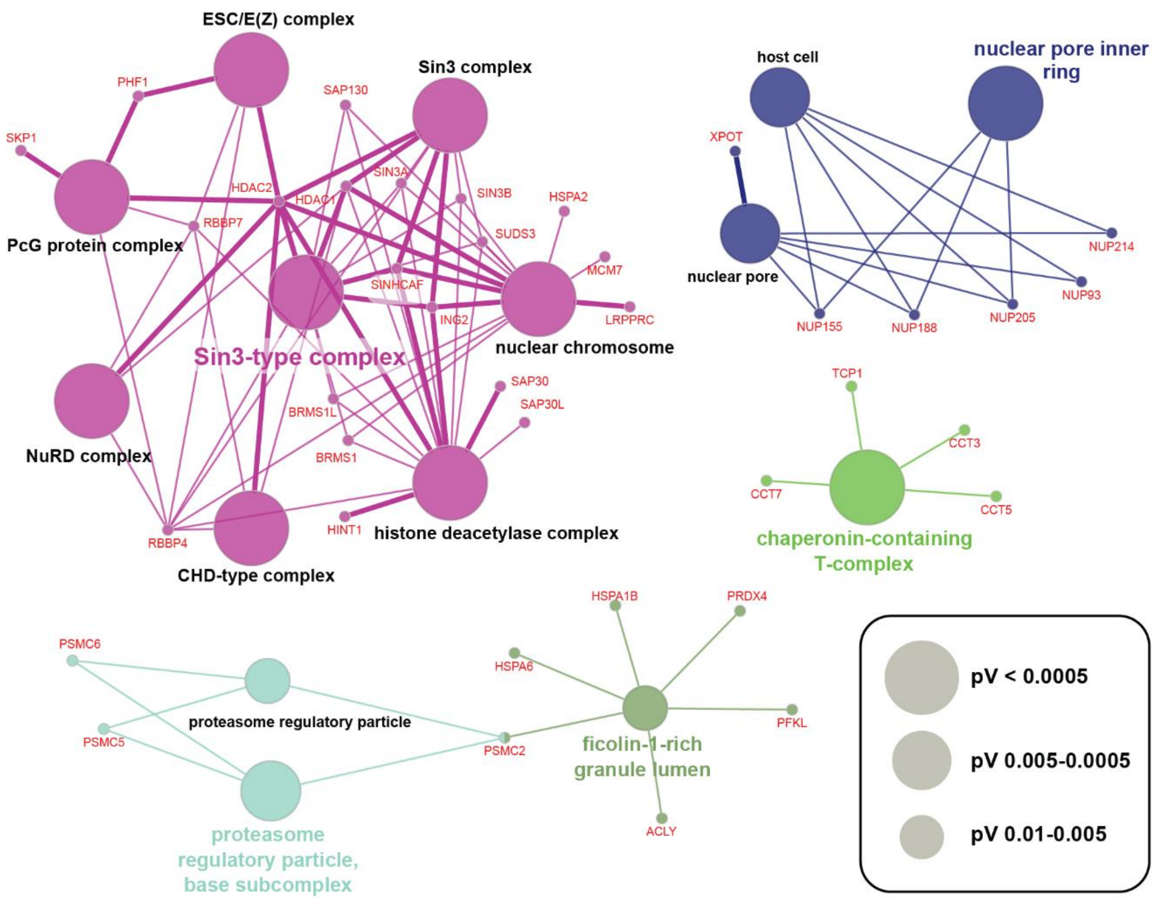


Figure S1. **Biological process associated with SAP25.** Gene Ontology terms (Biological process) for the proteins copurified with SAP25 Stably expressed in Flp-In™-293 stable cell line was determined using ClueGO with default setting.
